# Label-free high-speed wide-field imaging of single microtubules using interference reflection microscopy

**DOI:** 10.1101/273086

**Authors:** Mohammed Mahamdeh, Steve Simmert, Anna Luchniak, Erik Schäeffer, Jonathon Howard

## Abstract

When studying microtubules in vitro, label free imaging of single microtubules is necessary when the quantity of purified tubulin is too low for efficient fluorescent labeling or there is concern that labelling will disrupt its function. Commonly used techniques for observing unlabeled microtubules, such as video enhanced differential interference contrast, dark-field and more recently laser-based interferometric scattering microscopy, suffer from a number of drawbacks. The contrast of differential interference contrast images depends on the orientation of the microtubules, dark-field is highly sensitive to impurities and optical misalignments, and interferometric scattering has a limited field of view. In addition, all of these techniques require costly optical components such as Nomarski prisms, dark-field condensers, lasers and laser scanners. Here we show that single microtubules can be imaged at high speed and with high contrast using interference reflection microscopy without the aforementioned drawbacks. Interference reflection microscopy is simple to implement, requiring only the incorporation of a 50/50 mirror instead of a dichroic in a fluorescence microscope, and with appropriate microscope settings has similar signal-to-noise ratio to differential interference contrast and fluorescence. We demonstrated the utility of interference reflection microscopy by high speed imaging and tracking of dynamic microtubules at 100 frames per second. In conclusion, the image quality of interference reflection microscopy is similar to or exceeds that of all other techniques and, with minimal microscope modification, can be used to study the dynamics of unlabeled microtubules.

## INTRODUCTION

*In vitro* imaging of single microtubules has proved to be an important tool to study microtubule dynamics (Hotani and Horio, 1988; Walker et al., 1988), mechanics (Dogterom and Yurke, 1997; Gittes et al., 1993), and how they are regulated (Gell et al., 2010). Typically, fluorescence microscopy, in particular total internal reflection microscopy (TIRF), is used for imaging microtubule dynamics and gliding assays due to its high signal-to-background noise ratio (SBR, a measure of the optical contrast of the microtubule) and the wide range of available fluorescent labels (Gell et al. 2010). Efficient fluorescent labeling of tubulin requires a high tubulin concentration but often the yield of purified tubulin is too low (Widlund et al., 2012). In other cases, fluorescence-induced photo-damage is a concern (Vemu et al., 2016) especially if oxygen scavengers can react with the molecules of interest. In such cases, label free imaging is required.

There are many label-free imaging techniques available for imaging microtubules. Excellent techniques like video enhanced differential interference microscopy (DIC) and dark-field microscopy (DF) have been regularly used for almost 30 years (Bormuth et al., 2007; Gittes et al., 1993; Hotani and Horio, 1988; Walker et al., 1988). With DIC it is possible to image a large field of view at high frame rates. The drawback of using DIC is that the contrast of microtubules depends on their orientation with respect to the axis of the Nomarski prims. Although this can be solved (Shribak et al., 2008), the solution adds extra layers of complexity to the set up. Dark-field can visualize microtubule at high SBR assuming very clean solutions and surfaces. Otherwise, stray light caused by misalignment or light scattered by impurities will overwhelm the microtubule signal. Another technical inconvenience is that both techniques require the use of high numerical-aperture (NA) condensers that limit access to the sample during experiments. Other than DIC and dark-field, a number of techniques have emerged over the years. For example, Amos and Amos (Amos and Amos, 1991) imaged single microtubules by confocal reflection microscopy, requiring minimal modification to a confocal setup. Medina et al (Medina, 2010) visualized microtubules by defocusing the bright-field microscope, though the contrast was low even after image processing. Recently, Andrecka et al., used interferometric scattering (iSCAT) microscopy to image microtubule dynamics at high frame rates (Andrecka et al., 2016). iSCAT is capable of label-free imaging of single proteins but suffers from a limited field of view (practically on the order of 20x20 µm^2^) and requires a complex setup that involve lasers and laser scanners (Ortega Arroyo et al., 2016). Rotating-coherent-scattering microscopy (ROCS) which is also based on laser scattering was used to image microtubules far from the surface (Koch and Rohrbach, 2018). Just like iSCAT, ROSC requires using laser and scanners which complicates its implementation. Another recent technique uses spatial light interference microscopy (SLIM); while capable of imaging a large field of view (200x200 µm^2^) the frame rate is low (6.25 fps) (Kandel et al., 2017).

All the aforementioned techniques are capable of imaging single microtubules, but each fails to meet one or more of the following four criteria: (i) large field of view for higher throughput and to take advantage of the large chips of the new scientific CMOS cameras, (ii) high frame rate to improve temporal resolution, (iii) good signal-to-background noise ratio to achieve high spatial precision, and (iv) ease of implementation with low cost. In this work, we demonstrate that interference reflection microscopy (IRM) can image single label-free microtubules and satisfies all criteria. In IRM the image is formed by the interference between the light reflected from the glass-medium interface and that reflected from the medium-sample interface (Limozin and Sengupta, 2009; Weber, 2003). Since its first use in biology by Curtis in 1964 (Curtis, 1964), IRM, and the more advanced implementation known as reflective interference contrast microscopy, have been used intensively to probe cell adhesion to surfaces and other applications (Barr and Bunnell, 2001; Verschueren, 1985). Here we show that it is possible to image single microtubules using IRM in its simplest configuration.

## METHODS

IRM was implemented by adding a 50/50 mirror (Chroma, Bellows Falls, VT USA) to a Nikon inverted fluorescence microscope (Ti Eclipse, Nikon, Melville, NY USA) in the position where usually a dichroic mirror is placed. No additional excitation or emission filters are needed. The sample was illuminated by a Sola light engine (Lumencore, Beaverton, OR USA) attached to the microscope’s epi port. The NA of the illumination was set by the aperture iris. The field diaphragm was adjusted to optimize the contrast as presented in the Results section. The illumination light was partially reflected to the objective (100x/1.49 Apochromat, Nikon, Melville, NY USA) and the light reflected from the sample was collected by the same objective and projected –after passing through the 50/50 mirror– onto a 16 bit sCMOS camera (Zyla 4.2, 6.5 µm pixel size (65 nm at image plane), 2048x2048 pixel^2^ chip size, 72% quantum efficiency, Andor, Belfast, Scotland). The illumination was adjusted to nearly saturate the camera’s dynamic range. Experiments were performed in a flow channel formed by two parafilm strips sandwiched between 18x18 mm^2^ and 22x22 mm^2^ coverslips (#1.5H, Marienfeld, Germany) or between a 22x22 mm^2^ coverslip and a one inch slide (Thermo Scientific, Waltham, MA USA) (Gell et al., 2010). The coverslips were cleaned using piranha solution and rendered hydrophobic by silanization (Gell et al., 2010). In non-dynamic assays GMPCPP stabilized, TAMRA labeled (tetramethylrhodamine Ex: 550 nm, Em: 580 nm) bovine microtubules were imaged at room temperature. The microtubules were either passively attached to the hydrophobic surface or fixed by an anti-TAMRA antibody (Life Technologies, Waltham, MA USA). For dynamic assays, the same configuration was used and microtubule growth was initiated by flowing in polymerization solution (80 mM PIPES/KOH, pH 6.9, 1 mM ethylene glycol tetra-acetic acid (EGTA), 1 mM MgCl_2_, 2 mM GTP and 7.5 µM unlabeled bovine tubulin) at 34°C. Dynamic assays were imaged at slow (0.2 fps) and fast (100 fps) frame rates. In some dynamic assays the minus and plus end of the microtubules seeds were labeled using Alexa488 labeled (Ex: 490 nm, Em: 525 nm) tubulin. In all IRM imaging a long pass filter (LP 594 nm, Chroma, Bellows Falls, VT USA) was inserted into the illumination path to avoid exciting the fluorophores. Such a filter is unnecessary if unlabeled tubulin is used. No anti-fade reagents (Gell et al., 2010) were used.

To enhance contrast and eliminate illumination irregularities and static noise, a background image was subtracted from the acquired images. The background image was generated by averaging 32 or 100 images acquired before flowing in the microtubules or it was the median of a 100 images acquired while moving the sample at a high speed after imaging. Further enhancement of images included averaging, Fourier filtering or both (Bormuth et al., 2007). All image processing was performed using Fiji (Schindelin et al., 2012). Tracking of microtubules was done using the tracking software FIESTA (Ruhnow et al., 2011).

For DIC imaging of microtubules, a high NA oil immersion condenser equipped with a high resolution prism (NA = 1.4, Nikon, Melville, NY USA) was used for illuminating the sample. To enhance the contrast of the images, a bias retardation of one tenth of the wavelength was applied. Background images were generated by averaging 100 images taken 1-2 µm from the surface. As for IRM images, the images could be further enhanced by averaging and/or filtering.

Fluorescence images were obtained by TIRF microscopy. The microtubules were excited by a 561 nm laser at a power of ∼ 1 mW (at the sample plane) with an exposure time of 100 ms.

As a metric to assess IRM and compare it to different techniques we measured the signal-to-background noise ratio of a microtubule as defined by the average intensity of the microtubule signal (intensity of the microtubule minus the intensity of the background) divided by the standard deviation of the background. The background was defined in a region close to the microtubule, as described below.

For IRM and TIRF images, the signal was obtained by first isolating the microtubule in a rectangle region of interest (ROI) that was slightly longer than the microtubule along its axis and 60 pixels in the perpendicular direction. Next, the cross section profile of the microtubule was measured by taking a line scan perpendicular to the microtubule axis. The line width was set to equal the microtubule length. This way, every point on the cross section profile was an average of all pixels along the microtubule axis. Then, the signal was measured as the difference between the profile peak and the background. The background was measured by thresholding the microtubule image to separate the microtubule from the background and then averaging all pixels below the cut off intensity. In the case of DIC images, the signal-to-background noise ratio was determined as the peak to peak difference of the averaged signal divided by the standard deviation of the background noise (Bormuth et al., 2007).

We also measured the contrast sensitivity coefficient (CSC) define as the microtubule signal divided by the standard deviation of the microtubule signal. Here, the signal is the mean of all the pixels’ values above the threshold cut off after background subtraction. The noise is the standard deviation of the signal. CSC measurement is applicable to IRM and TIRF microtubule images but not DIC because the standard deviation along the microtubule will be higher than the signal. This is because the signal will nearly vanish since the intensities above and below the background tend to cancel each other.

## RESULTS AND DISCUSSION

Microtubules are readily visible in IRM without background subtraction. The microtubules appeared dark against a brighter background (Fig, 2a). Reducing the field of view size by the field diaphragm reduced the amount of stray light reaching the camera and improved the contrast of the microtubule (Fig. 1a). We found that opening the field diaphragm up to 70% of the field of view was a good balance between improving contrast and maintaining a large imaging area. In addition to the field diaphragm, the contrast depended on the illumination NA (Weber, 2003). Qualitatively, without background subtraction, the microtubules visibility was highest in the NA range 0.7-1.1 (Fig. 1b). This agreed with SBR measurements of background-subtracted images as a function of illumination NA (Fig. 1c). The SBR of a single frame of background-subtracted images was 6.8 ± 0.8 (mean ± SD, *N* = 41). To further improve the SBR, the images were either averaged or filtered or both. Upon averaging, the SBR initially increased with a power law of 0.25, less than the 0.5 dependence expected for a photon-shot-noise limited image. This deviation can be explained by the presence of other sources of noise such as dirt and drift. As the number of averaged frames increased, the SBR saturated as the noise in the image was no further reduced by averaging (supplementary Fig, 2a). By careful examination of the noise, we found that as the number of averaged frames approached the number of averaged background images the gained noise reduction saturated (supplementary Fig, 2b). The average of 10 images, increased the SBR to 11.6 ± 1.6. A comparable increase to 11.2 ± 1.2 was obtained by using a low-pass Fourier filter. Filtering the averaged images results in a final SBR of 17.4 ± 2.4 (Fig, 2).

**Fig 1.**
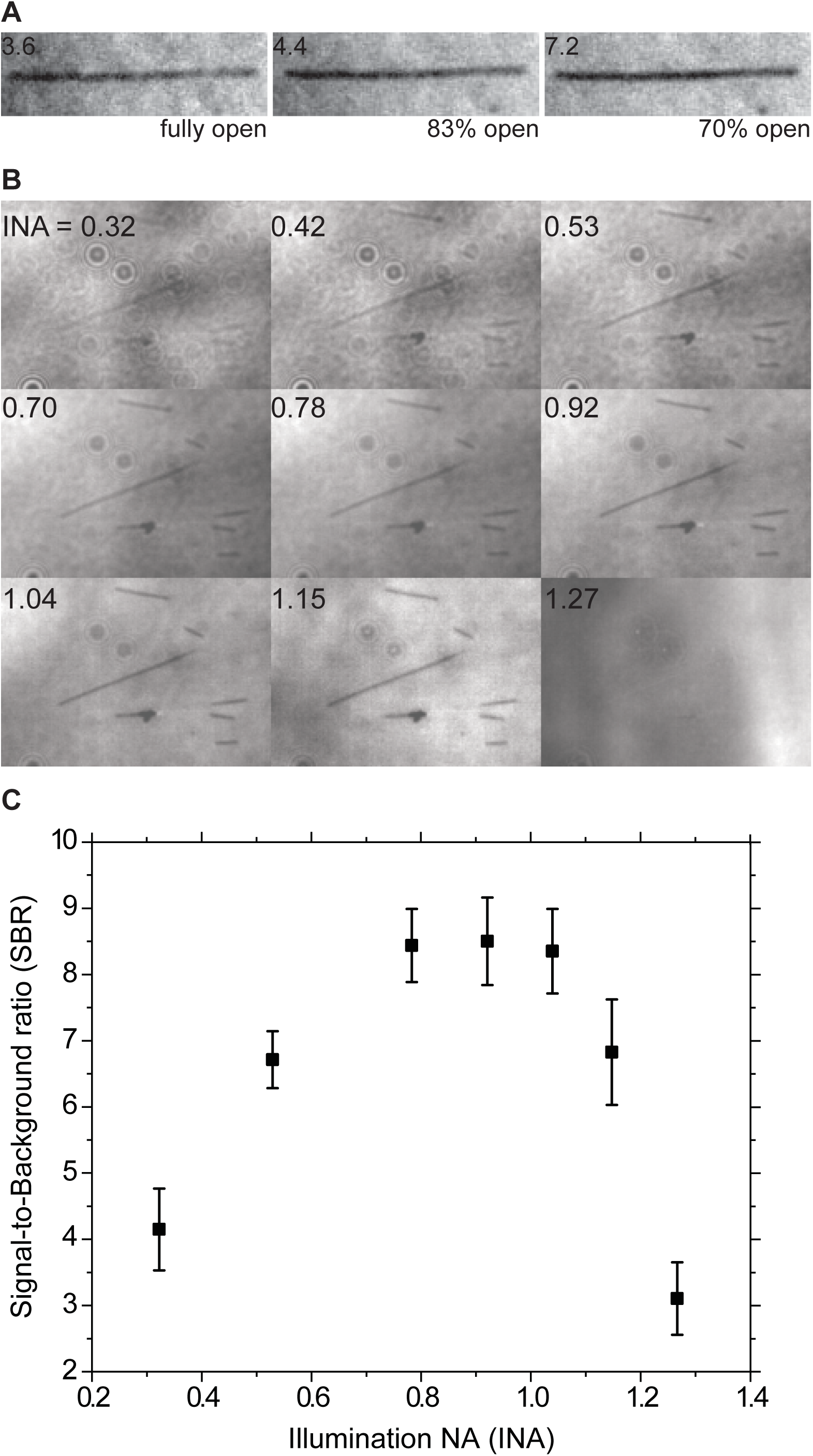

**Fig 2.**
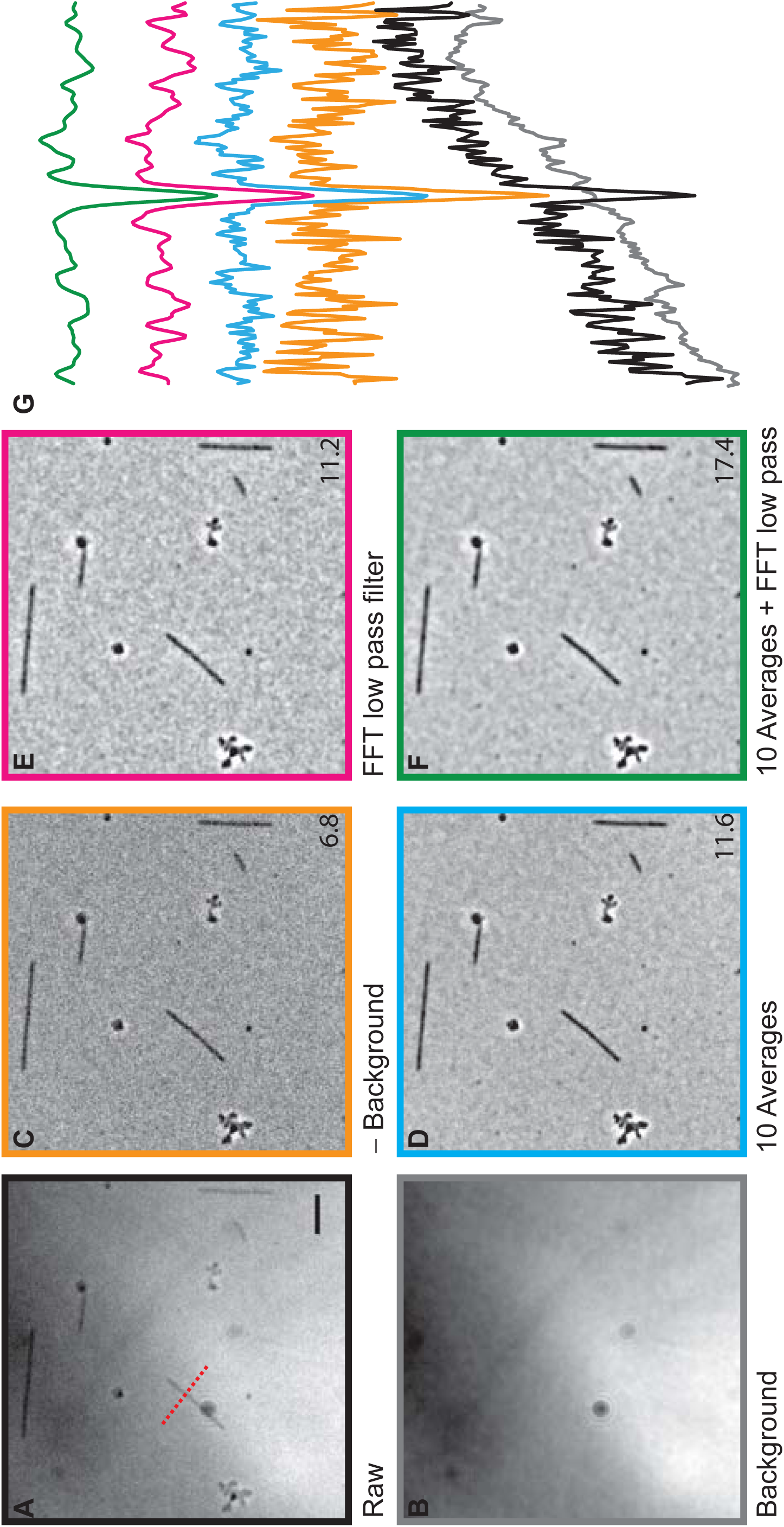

To compare the performance of IRM to other techniques, the same field of view of microtubules was imaged by IRM, TIRF and DIC and the SBR for each image was measured (Fig, 3a). For TIRF of highly labelled microtubules (30%, to reduce speckles (Waterman-Storer et al., 1998)), the SBR was 40 ± 10 (mean ± SD, *N* = 41) prior to photobleaching. In terms of CSC, both IRM and TIRF images were similar, CSC_TIRF_= 2.64 ± 0.13 (mean ± SD) compared to CSC_IRM_= 2.7 ± 0.18. DIC images had a lower SBR (10 ± 2) than IRM. Aside from the higher SBR, the advantage of using IRM over DIC is that the contrast is independent of microtubule orientation as can be clearly seen in Fig, 3a. Such dependence has an impact when tracking microtubules imaged in DIC (Danuser et al., 2000; Janson and Dogterom, 2004).

**Fig 3.**
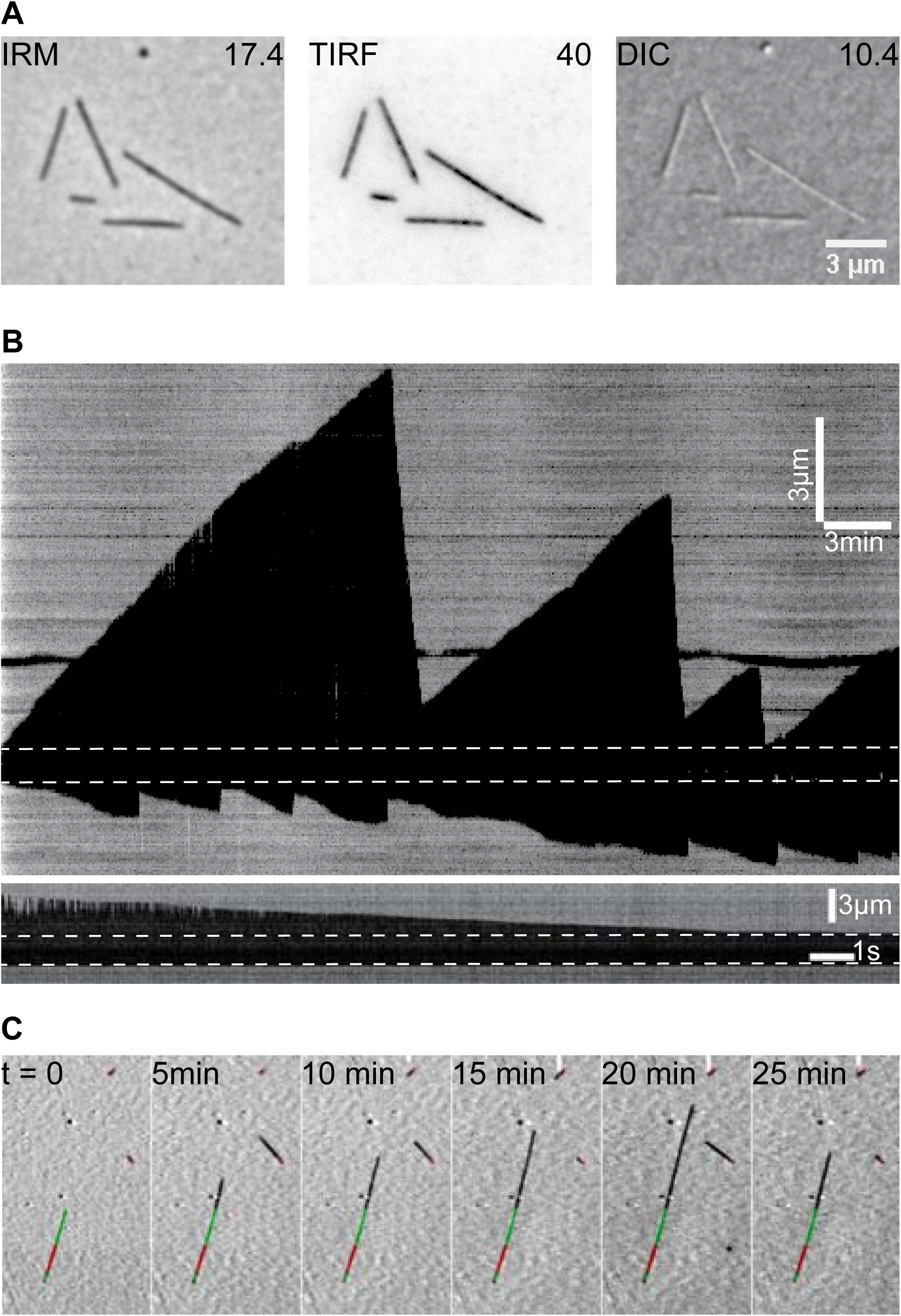

To demonstrate the capability of IRM, dynamic microtubules were imaged. Averaged background-subtracted images were taken at a frame rate of 0.2 fps. Growth and shrinkage phases where easily observed in the videos and kymographs (Fig, 3b, Supplementary video 1). While such low frame rate is suitable for measuring growth rate and catastrophe frequency, it is more challenging to measure the shrinkage rates which are an order of magnitude higher than growth rates. For this reason, shrinking microtubules were imaged at 100 fps which provided a detailed image of the shrinking end (Fig, 3b). At both frame rates, it was possible to track the microtubule ends using the tracking software FIESTA with a length precision of ≈20 nm (Supplementary video 1). In addition, IRM can be easily combined with fluorescence microscopy, thus freeing one of the fluorescence channels to image, for example, molecular motors, MAPs or specially modified tubulins (Fig, 3c).

In TIRF and DIC, thermal fluctuations of long microtubules away from the surface lead to loss of contrast. In IRM, such fluctuations lead to a reverse in microtubule contrast, which can be used to measure the height of the microtubule, similar to fluorescence interference contrast microscopy (Kerssemakers et al., 2006; Lambacher and Fromherz, 1996) (supplementary video 2). Recently, the ability to measure distances from the surface using IRM was used as a mean to calibrate the TIRF evanescent field in a combined IRM-TIRF-optical tweezers setup using a light emitting diode for illumination (Simmert et al., 2018). It is also worth mentioning that IRM is flexible in terms of what objective to use: it was possible to image microtubules using high end and low end objectives as well as phase and DIC objectives. The presence of a Nomarski prism or the phase ring did not seem to influence the images.

The main drawback of IRM is its sensitivity to drift which is noticeable when imaging for extended periods of time (Supplementary Fig, 3). This requires the setup to be thermally stabilized (Mahamdeh and Schäffer, 2009) or the image to be drift corrected (Carter et al., 2007; Kim and Saleh, 2008; Ortega Arroyo et al., 2016).

In conclusion, IRM proved to be a powerful tool for imaging unlabeled microtubules in surface assays. It only requires a minor, cost effective modification to any epi-microscope, requires a onetime alignment, it is capable of high-speed wide-field (100 fps at full chip) imaging of microtubules at a high spatial precision for long periods of time and can be combined easily with other techniques.

